# Effect of PAPE Induced by Different Squat Loads on Jump Performance in Collegiate Female Basketball Players

**DOI:** 10.1101/2025.11.06.687070

**Authors:** Xu Si, Yu Hang Liu, Xin Ru Zhou, Shao Jie Ning, Qing Bo Li

## Abstract

**Background:** Post-activation performance enhancement (PAPE) can acutely augment explosive performance, yet the optimal prescription of squat load and recovery for female basketball players remains unclear.

**Methods:** Twenty-eight collegiate women’s basketball athletes were randomly allocated to 90%, 80%, or 70% of one-repetition maximum (1RM) back-squat groups, or a control group. Athletes completed 3×3 parallel back squats, after which countermovement jump (CMJ), single-leg and double-leg approach-jump heights were assessed at baseline and 4, 8, and 12 min. CMJ kinetics—peak power output, vertical ground-reaction force, and flight time—and lower-limb surface EMG were recorded concurrently.

**Results:** Loads ≥80%1RM reliably elicited PAPE, with improvements demonstrating clear load-dependence and time specificity. Double-leg approach-jump height peaked at 8 min in the 90%1RM group (40.50 ± 1.73 cm). Single-leg approach-jump height was maximized at 8 min in the 80%1RM group (40.00 ± 0.82 cm), exceeding the control condition. Kinetic and EMG analyses indicated that 90%1RM produced a delayed rectus femoris activation peak (8 min) and a later CMJ power peak (12 min), whereas 80%1RM facilitated earlier gastrocnemius activation (4 min) with a stable output profile; performance benefits with 70%1RM were minimal.

**Conclusions:** A moderately high load (80%1RM) paired with an 8–12-min recovery window strikes a favorable balance between neural potentiation and fatigue, offering a practical pre-competition activation strategy to enhance anaerobic power and jump performance in female basketball players.

## 1. introduction

Basketball demands exceptional explosive power, speed, and agility^[1]^. As the pace of modern play accelerates, jumping actions—rebounding, blocking, and a variety of jump shots—have become decisive factors in game outcomes^[2]^. Vertical jump capacity not only confers direct scoring and defensive advantages but also reflects overall competitive proficiency^[3]^. Female athletes, however, often encounter unique challenges in explosive performance owing to sex-related differences in neuromuscular control, endocrine milieu, and muscle morphology^[4]^ ^[5]^ ^[6]^. Identifying acute, safe, and effective strategies to boost jump performance in women’s basketball is therefore of immediate practical relevance for pre-competition preparation and performance optimization^[7]^.

Post-activation performance enhancement (PAPE) denotes the transient augmentation of strength, power, and rate of force development following a high-intensity conditioning activity, with effects lasting from minutes to as long as 48 hours^[8]^. Mechanistically, PAPE is attributed primarily to increased Ca²⁺ sensitivity via phosphorylation of the myosin regulatory light chain, alongside heightened motor-unit recruitment, more efficient neural transmission, and improved force transfer associated with reduced muscle pennation angles^[9]^ ^[10]^. Its magnitude and time course are shaped by both individual and programming factors: greater strength and training experience and a higher proportion of type II fibers tend to amplify responses, whereas aging may blunt them; sex and phase-specific physiology can further modulate outcomes^[11]^ ^[12]^. On the programming side, the induction mode, load–repetition configuration, and recovery interval jointly determine the point at which fatigue dissipates and potentiation emerges^[13]^ ^[14]^ ^[15]^。

Parallel back squat (PS)—defined here as squatting to a depth at which the superior thigh is parallel to the floor—is not only foundational for long-term strength development but also a potent neuromuscular primer^[16]^.Esformes and Bampouras ^[17]^ suggested that deeper PS may more readily elicit PAPE, potentially via greater activation of hip extensors (e.g., gluteus maximus) and consequent improvements in lower-limb force output. Manipulating PS intensity meaningfully alters muscle-activation profiles: higher loads (>85% 1RM) favor recruitment of high-threshold motor units and neural adaptations, whereas moderate loads (60–80% 1RM) capitalize on velocity–force coupling to translate more directly into rapid force production^[18]^ ^[19]^. This load-dependent responsiveness positions loaded squats as a practical intervention for acutely enhancing vertical-jump performance through PAPE^[20]^ ^[21]^.Nevertheless, the interplay between load magnitude and recovery interval—and potential sex-specific responses linked to women’s neuromuscular characteristics—remains insufficiently defined.

Therefore we compared three PS intensities (70%, 80%, 90% 1RM) against an unloaded control to delineate the acute intensity–time profile of PAPE in female basketball players. Performance was assessed at 4-, 8-, and 12-minutes post-intervention relative to baseline. During countermovement jumps (CMJ), vertical ground-reaction forces and lower-limb surface EMG were recorded to interrogate mechanical and neuromuscular mechanisms. CMJ outcomes included peak power, flight time, and ground-reaction-force metrics, along with RMS EMG of targeted muscles; single- and double-leg approach-jump reach heights were concurrently measured to capture functional jumping performance. We hypothesized that all load conditions (70%, 80%, 90% 1RM) would outperform control (main PAPE effect), and that load × time interactions would be evident: specifically, 80% 1RM would yield the greatest improvement at 4 minutes, whereas potentiation with 90% 1RM would become more prominent at 8–12 minutes, matching or surpassing the 80% 1RM condition.

## 2 methods

### 2.1 Participants

To ensure adequate statistical power, an a priori sample size calculation was performed using G*Power v3.1.9.2 (Düsseldorf, Germany). The planned analysis was a four-group mixed-design repeated-measures ANOVA (between-subjects factor: group; within-subjects factor: time points = 3). Based on prior literature, we set the effect size at f = 0.25 (approximately d = 0.5), power at 0.80, and α = 0.05; the required total sample size was 28 participants (7 per group). We ultimately enrolled 28 collegiate female basketball players, all of whom met the training-level classification of McKay et al.^[22]^ (Tier 3; see Table 1). Inclusion criteria were: (a) ≥6 years of systematic basketball training; (b) resistance training 2–3 times per week during the past 6 years; (c) no lower-limb injury or other condition affecting training or testing for ≥6 months prior to the experiment; (d) test sessions scheduled outside of menstruation; and (e) no history of smoking or alcohol abuse. Participants abstained from resistance (strength) training for 48 h before testing and maintained regular dietary habits for 2 weeks before the experiment, avoiding creatine and any ergogenic supplements or stimulants; baseline anthropometrics (e.g., height, body mass) were recorded at the start of testing. All participants provided written informed consent and were explicitly informed that they could withdraw at any time without penalty; to minimize bias, the specific study objectives were not disclosed until after testing. Participant recruitment took place from February 18, 2025, to June 30, 2025. The study protocol was approved by the Ethics Committee of Tianjin University of Sport (Approval No.: TUS2025-103) and conducted in accordance with the Declaration of Helsinki.

**Table 1.** Participant Characteristics (Mean ± SD)

| Subject | 1RM90% Group | 1RM80% Group | 1RM70% Group | Control Group | P-value |
| --- | --- | --- | --- | --- | --- |
| Age (years) | 20.50 ± 0.57 | 20.25 ± 1.5 | 21.5 ± 2.38 | 20.32 ± 0.14 | P=0.106 |
| Height (cm) | 167.25 ± 2.21 | 168.25 ± 3.09 | 168.30 ± 2.23 | 170.02 ± 1.43 | P=0.602 |
| Body Mass (kg) | 56.50 ± 4.20 | 56.75 ± 9.56 | 58.50 ± 4.50 | 59.25 ± 1.86 | P=0.507 |
| Training Age (years) | 3.23 ±0.76 | 4.67 ±0.48 | 3.32 ±0.63 | 3.76 ±0.58 | P=0.577 |
| 1Repetition Maximum Squat | 99.57 ± 8.11 | 98.80 ± 6.06 | 98.30 ± 5.36 | 97.01 ± 4.10 | P=0.153 |

### 2.2 Experimental Design

This study adopted a randomized, parallel group controlled design. Twenty-eight female basketball players meeting the inclusion criteria were allocated by a random-number table to four groups: 1RM90% (n = 7), 1RM80% (n = 7), 1RM70% (n = 7), and a control group (n = 7). The aim was to compare the acute effects of different squat loads on countermovement jump (CMJ), single-leg approach jump reach (SLAJ), and double-leg approach jump reach (DLAJ), and to examine their trajectories at post-intervention time points (4, 8, and 12 min). The loading prescription—the experimental independent variable—was standardized as follows: in the load groups, participants performed barbell back squats for 3 sets × 3 repetitions, with 1 min 30 s inter-set rest, to a depth where the thighs were parallel to the floor; the control group received no loading stimulus and only underwent the same measurement schedule. Intensities were set at 70%, 80%, and 90% of 1RM, and CMJ, SLAJ, and DLAJ were re-assessed at 4, 8, and 12 min after the intervention to delineate the acute time course. The intensity and prescription were based on prior evidence indicating that 50–95% 1RM can elicit post-activation performance gains^[23]^, with 70–90% 1RM offering a favorable balance between potentiation and muscle-damage risk^[24]^, and a 2–14 min window commonly used to observe the PAP–fatigue interplay^[25]^. Primary dependent variables included CMJ height/power and SLAJ/DLAJ reach height. For neuromuscular activation, surface EMG was collected only during CMJ, with MVC calibration for RMS normalization; SLAJ and DLAJ captured jump height only, without EMG or MVC. This decision reflects the substantial motion artifacts and inter-individual rhythm variability inherent to approach and arm-swing actions in SLAJ/DLAJ, as well as the potential for electrodes/leads to disrupt natural coordination. Accordingly, height was treated as the primary performance index for SLAJ/DLAJ, whereas neuromuscular activation mechanisms were characterized via the more standardized CMJ protocol.

### 2.3 procedure

The experiment was conducted from May to June 2025 in Laboratory Building A of the Human Performance Science facilities at Tianjin University of Sport. All testing sessions were scheduled on Tuesdays and Thursdays between 13:00 and 17:00 to control for environmental conditions and potential circadian influences. All procedures w ere administered by three strength-and-conditioning graduate students who had received standardized training, ensuring procedural consistency and data reliability. At the initial laboratory visit, we collected anthropometric measures (including body composition assessed with a Tanita MC-780MA, Japan). Participants then completed familiarization with the test movements and equipment. The experimental workflow comprised four phases: (1) preparation and warm-up; (2) baseline testing (on the baseline day only countermovement jump [CMJ], single-leg approach jump reach [SLAJ], double-leg approach jump reach [DLAJ], and surface EMG calibration were performed; the 1RM parallel back squat [SP] had been conducted three days prior to formal testing following a standardized protocol); (3) SP conditioning intervention (executed per group-specific prescriptions; the control group undertook quiet seated rest during the equivalent time window to match time and attentional demands); and (4) post-intervention assessments initiated at t = 4, t = 8, and t = 12 minutes (see Figure 1).

**Figure 1.**
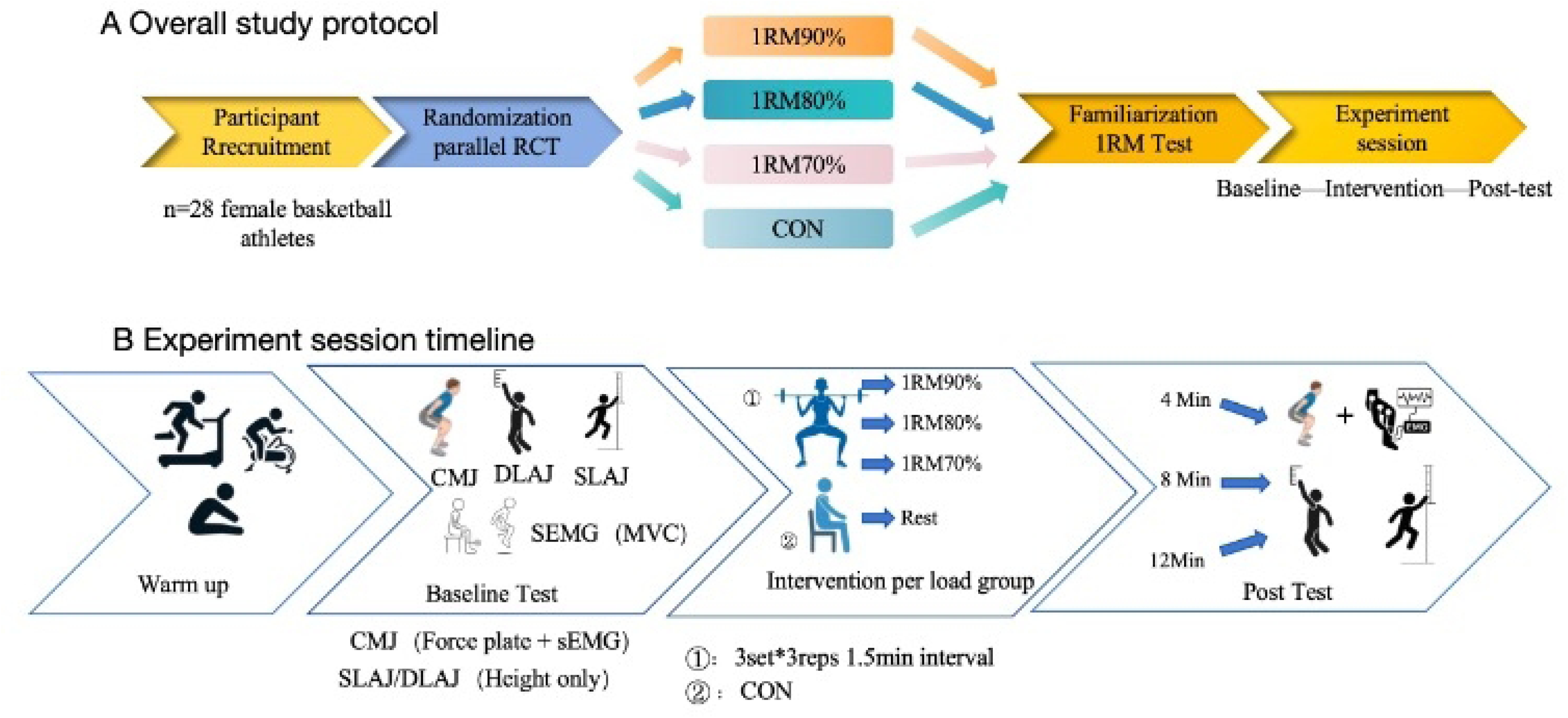
Panel A demonstration the overall study timeline; Panel B shows the detail timeline for the Experiment session. SLAJ, Single-Leg Approach Jump; DLAJ, Double-Leg Approach Jump; EMG, Surface Electromyography.

#### 1RM Parallel Back Squat (SP) Test

The 1RM SP test was completed three days prior to the formal intervention and scheduled within the same daily time window (13:00–17:00) to minimize circadian influences^[26]^。Participants wore standardized training shoes and used a 20-kg Olympic bar bell with calibrated plates. The test was conducted in a power rack with safety pins s et; one spotter was positioned on each side. The session was terminated immediately i f technical criteria were not met or any safety concern arose. The squat variation was fixed as a high-bar back squat; stance width was self-selected but had to remain con sistent in subsequent sessions. Squat depth was standardized to thighs parallel to the floor, judged from the side view by two assessors, with concordant ratings required for a valid lift. The eccentric phase was controlled at 2–3 s with no pause; the concentric phase was performed as fast as possible. No bouncing or compensatory swinging w as permitted. Warm-up and load progression followed the NSCA standardized protocol^[27]^: perform 5–10 warm-up repetitions at a moderate load, rest 1 min; increase the ini tial load by 10–20% for 3–5 repetitions, rest 2 min; increase by a further 10–20% for 2–3 repetitions, rest 2–4 min. During formal 1RM attempts, increase the load by 5–1 0% each attempt (adjusting to 2.5–5% near maximal efforts), with 2–4 min rest between attempts; after a failed attempt, decrease the load by 2.5–10% before retrying. The 1RM was determined within no more than five formal attempts. A successful 1RM required meeting the depth criterion, lifting without assistance, and exhibiting no obvious technical imbalance.

#### CMJ Test

Countermovement jump (CMJ) procedures followed the standardized protocol in th e Swiss Olympic Fitness Testing Manual [22]. CMJ was assessed using a Kistler force platform (9257B, Switzerland), with a 5-s recording window per trial and a sampling frequency of 2000 Hz. Prior to testing, participants completed a full warm-up (dyna mic stretching and low-intensity jumping). The laboratory environment was kept quiet and free from distractions, and the force plate was calibrated before data collection to ensure accuracy. Participants performed CMJs with hands on hips to eliminate arm-swing effects. From an upright stance, they executed a rapid countermovement to approximately 90° knee flexion and immediately jumped maximally. Throughout the movement, athletes were instructed to maintain trunk stability, avoid anterior–posterior sway a nd knee valgus, achieve full lower-limb extension with simultaneous take-off, and land stably near the center of the force plate.The following variables were selected for analysis: jump height, relative peak power output, and maximal vertical ground-reaction force. Jump height and peak power were computed from flight time and the force–time signal recorded by the platform. Data were processed in BioWare software (version 5.3.0.7); time resolution was set to 1/1000 s (1 ms). Each participant performed 3–5 tri als with 1–2 min inter-trial rest to ensure adequate recovery; the best trial was retained for statistical analyses. Trials showing technical compensation or anomalous signals were corrected when possible and otherwise discarded to ensure data reliability.

#### Single- and Double-Leg Approach Jumps (SLAJ/DLAJ)

SLAJ and DLAJ were assessed using a Vertec vertical jump device (resolution ≥ 0.5 cm). Standing reach height was recorded as baseline before testing. Each task comprised three valid attempts with 60–90 s inter-trial rest; jump height was calculated as highest reach − standing reach. For SLAJ, participants took off from the dominant leg and touched the vanes with the ipsilateral hand. For DLAJ, athletes executed a “step-through–gather step” over the final two strides, performed a simultaneous two-foot t ake-off, and touched the vanes with the dominant hand. Before the first attempt, each athlete determined—then maintained across trials—their approach stride count, start li ne, and take-off zone, which were clearly marked with tape (any change rendered the attempt invalid and required a re-test after equivalent recovery). Natural arm swing a nd individual rhythm were permitted; no double take-offs, double touches, or intentional sliding were allowed. Attempts were deemed invalid and repeated if any of the foll owing occurred: obvious disruption of rhythm or balance at take-off, insufficient knee/ hip extension at take-off, contact with a non-valid portion of the vanes, or unclear va ne contact. Two recorders independently read the measurement and compared values immediately; if the discrepancy exceeded 0.5 cm, a third adjudication was used. If the difference between the two best scores for the same task exceeded 2 cm, one additional re-test was performed after equivalent recovery.

#### Surface EMG acquisition and calibration

Surface EMG was collected only during CMJ to evaluate neuromuscular activation, using a NORAXON system (USA) at 1500 Hz. Following SENIAM recommendations^[28]^, the skin was prepared (alcohol cleansing → shaving → light abrasion) before applying disposable Ag/AgCl electrodes; electrodes were aligned with the muscle-fiber direction over the muscle belly with ∼20 mm center-to-center spacing, and a reference electrode was placed on the ipsilateral superior border of the patella (bony landmark) to reduce noise. Three target muscles and electrode placements were as follows: rectus femoris—anterior thigh at the midpoint of the line connecting the anterior superior iliac spine and the superior border of the patella; biceps femoris long head—posterolateral thigh near the proximal half of the line connecting the ischial tuberosity and fibular head; lateral gastrocnemius—proximal one-third of the most prominent muscle belly on the lateral shank along the line from the fibular head to the superior border of th e Achilles tendon. Prior to data collection, maximal voluntary contractions (MVCs) we re recorded for each muscle to obtain RMS_MVC: rectus femoris—seated, knee at 60° flexion, resisted knee extension for 3 s; biceps femoris long head—prone, knee at 30° flexion, resisted knee flexion for 3 s; lateral gastrocnemius—seated, knee at 90° flex ion, resisted plantarflexion for 3 s; with 2 min rest between MVCs. Signals were processed in MR3 (v3.18.136) with full-wave rectification, 20–500 Hz band-pass filtering, and smoothing. For CMJ analyses, the concentric phase window extended from the on set of the countermovement (i.e., when the body’s center of mass began to accelerate downward) to toe-off; within this window, the RMS of each EMG trace was computed and then normalized to the corresponding muscle’s MVC RMS to yield %MVC. Processed data were exported for statistical analysis.

### 2.4 Statistical analyses

Between-group comparisons of demographic variables were conducted using one-w ay ANOVA; pairwise comparisons employed independent-samples t tests with Bonferroni adjustment for multiple testing. Normality was first assessed with the Shapiro–Wilk test; if normality or homoscedasticity was violated, demographic variables were analy zed with the Kruskal–Wallis test (or the Mann–Whitney U test for two-group compari sons), and nonparametric alternatives were applied to repeated-measures data as needed. Primary effects were tested via a mixed-design repeated-measures ANOVA with group (four levels: loading intensities/control) as the between-subjects factor and time (4, 8, 12 min) as the within-subjects factor, i.e., a 4 (group) × 3 (time) design. Mauchly’s test was used to examine sphericity; when violated, Greenhouse–Geisser corrections w ere applied. Statistical significance was set at α = 0.05 (two-tailed). When significant main or interaction effects were detected, simple-effects analyses were performed follo wed by Bonferroni-adjusted post hoc comparisons. Effect sizes were reported as η² an d interpreted using the following thresholds: η² = 0.01 (small), η² = 0.06 (medium), a nd η² > 0.14 (large)^[29]^。Data management and descriptive statistics were performed in Microsoft Excel 2019, and inferential analyses and visualizations were conducted in GraphPad Prism 10 and SPSS 26.0; results are presented as mean ± SD.

## 3 Result

A 4 (group: 90%1RM, 80%1RM, 70%1RM, control) × 4 (time: baseline, 4, 8, 12 min) mixed-design repeated-measures ANOVA was employed. Sphericity for time main effects and interactions was examined with Mauchly’s test; when violated, Greenhouse–Geisser corrections were applied. Effect sizes are reported as partial eta squared (ηp²). Statistical significance was set at α = 0.05 (two-tailed), and post hoc comparisons were adjusted using the Holm–Bonferroni procedure.

For DLAJ, there were significant main effects of group (F(3, 24) = 7.29, p = 0.002, ηp² = 0.31) and time (F(2.25, 54.0) = 19.85, p < 0.001, ηp² = 0.55), as well as a significant group × time interaction (F(6.75, 54.0) = 5.64, p = 0.004, ηp² = 0.26) (Figure 2A). Simple-effects analyses showed that at 4 min the 80%1RM group (32.50 ± 1.30 cm) was significantly lower than all other groups (p < 0.05). At 8 min, all loading groups outperformed control (control: 37.18 ± 0.58 cm; 90%1RM: 40.50 ± 1.73 cm; 80%1RM: 39.75 ± 1.25 cm; 70%1RM: 39.80 ± 1.50 cm; p < 0.05). These findings suggest that loading interventions may induce short-term enhancements in jump performance (Table 2).

**Figure 2.**
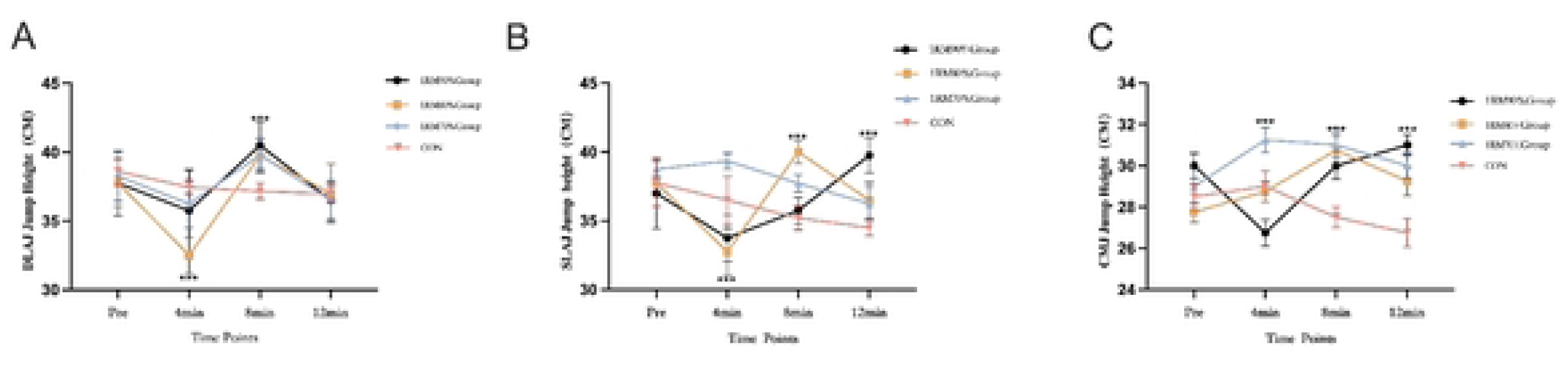
Time-course of jump height across recovery following the squat conditioning activity. A) Double-leg approach jump (DLAJ) height; B) Single-leg countermovement jump (SLCMJ) height; C) Countermovement jump (CMJ) height. Lines denote group means for the 90%1RM, 80%1RM, 70%1RM and control conditions at Baseline, 4 min, 8 min and 12 min post-intervention; error bars represent SD (n=7 per group) *P<0.05,**P<0.01, *** P<0.001

**Table 2.** Jump Height Under Different Conditioning Loads and Recovery Times (cm, Mean ± SD)

| Jump Type | Group | Baseline | 4 min | 8 min | 12 min |
| --- | --- | --- | --- | --- | --- |
| CMJ | 90%1RM | 30.00 ± 2.31 | 26.75 ± 1.89 <sup>a</sup> | 30.00 ± 3.37 | 31.00 ± 2.45 * # |
|  | 80%1RM | 27.75 ± 2.63 | 28.75 ± 0.96 | 30.75 ± 2.22 * # | 29.25 ± 1.50 |
|  | 70%1RM | 29.00 ± 3.27 | 31.25 ± 2.06 # | 31.00 ± 2.16 * # | 30.00 ± 2.00 |
|  | CON | 28.50 ± 0.58 | 29.00 ± 1.83 | 27.50 ± 1.29 | 26.75 ± 1.89 |
| DLAJ | 90%1RM | 37.50 ± 2.65 | 35.50 ± 3.11 | 40.50 ± 1.73 * # | 36.50 ± 1.29 |
|  | 80%1RM | 37.75 ± 1.71 | 32.50 ± 1.30 <sup>b</sup> # | 39.75 ± 1.25 * # | 37.00 ± 1.26 |
|  | 70%1RM | 38.25 ± 2.06 | 36.25 ± 1.85 | 39.80 ± 1.50 * # | 36.50 ± 1.70 |
|  | CON | 38.60 ± 0.93 | 37.45 ± 0.65 | 37.18 ± 0.58 | 36.95 ± 0.56 |
| SLAJ | 90%1RM | 37.00 ± 2.58 | 33.75 ± 1.70 | 35.75 ± 0.98 | 39.75 ± 1.25 * # |
|  | 80%1RM | 37.75 ± 1.71 | 32.75 ± 1.71 <sup>c</sup> | 40.00 ± 0.82 ** # | 36.50 ± 1.29 |
|  | 70%1RM | 38.80 ± 0.60 | 38.90 ± 0.30 # | 38.30 ± 1.00 # | 36.40 ± 1.10 |
|  | CON | 37.75 ± 1.71 | 36.50 ± 1.73 | 35.20 ± 0.86 | 34.50 ± 0.58 |

**Note:** CMJ = Countermovement Jump; DLAJ = Double-Leg Approach Jump; SLAJ = Single-Leg Approach Jump; CON = Control Group. Superscript # indicates the highest value (peak performance) within the same jump type and group across time. Significance symbols indicate a significant difference compared to the CON group at the same time point: \* $p < 0.05$ , \*\* $p < 0.01$ . Superscript letters indicate significant differences between experimental groups at the same time point: <sup>a</sup> Significantly lower than the 70%1RM group; <sup>b</sup> Significantly lower than all other groups; <sup>c</sup> Significantly lower than the 70%1RM group.

For SLAJ, significant main effects were observed for group (F(3, 24) = 9.13, p = 0.001, ηp² = 0.36) and time (F(2.25, 54.0) = 22.85, p < 0.001, ηp² = 0.59), along with a significant interaction (F(6.75, 54.0) = 4.79, p = 0.008, ηp² = 0.23) (Figure 2B). At 4 min, the 80%1RM group (32.75 ± 1.71 cm) was significantly lower than the 70%1RM group (p = 0.031). At 8 min, the 80%1RM group (40.00 ± 0.82 cm) was significantly higher than control (p < 0.01). At 12 min, the 90%1RM group (39.75 ± 1.25 cm) exceeded control (p = 0.022). Overall, the 80%1RM group peaked at 8 min, the 90%1RM group peaked at 12 min, the 70%1RM group showed a slight decline by 12 min, and control decreased throughout (Table 2).

For CMJ, there were significant main effects of group (F(3, 24) = 6.29, p = 0.002, ηp² = 0.31) and time (F(2.25, 54.0) = 18.85, p < 0.001, ηp² = 0.55), together with a significant interaction (F(6.75, 54.0) = 4.64, p = 0.004, ηp² = 0.26) (Figure 2C). At 4 min, the 70%1RM group (31.25 ± 2.06 cm) was significantly higher than the 90%1RM group (26.75 ± 1.89 cm; p = 0.017). At 8 min, both the 80%1RM and 70%1RM groups (30.75 ± 2.22 cm and 31.00 ± 2.16 cm, respectively) were significantly higher than control (27.50 ± 1.29 cm; p < 0.05). At 12 min, the 90%1RM group (31.00 ± 2.45 cm) was significantly higher than control (p = 0.026).

Relative peak power (PPO). There were significant main effects of group (F(3, 24) = 7.22, p = 0.002, ηp² = 0.36) and time (F(2.25, 54.0) = 22.85, p < 0.001, ηp² = 0.55), as well as a significant group × time interaction (F(6.75, 54.0) = 5.99, p = 0.001, ηp² = 0.27) (Figure 3A). At 4 min, the 90%1RM group (34.30 ± 3.76 W·kg⁻¹) was significantly lower than the 80%1RM and 70%1RM groups (p < 0.05). At 8 min, the 80%1RM (42.10 ± 2.15 W·kg⁻¹) and 70%1RM (41.30 ± 1.31 W·kg⁻¹) groups were significantly higher than control (p < 0.05). At 12 min, the 90%1RM (43.30 ± 5.18 W·kg⁻¹) and 80%1RM (42.90 ± 1.87 W·kg⁻¹) groups exceeded the other two groups (p < 0.05). Overall, higher loads exhibited stronger power output in the later recovery phase (Table 3).

**Figure 3.**
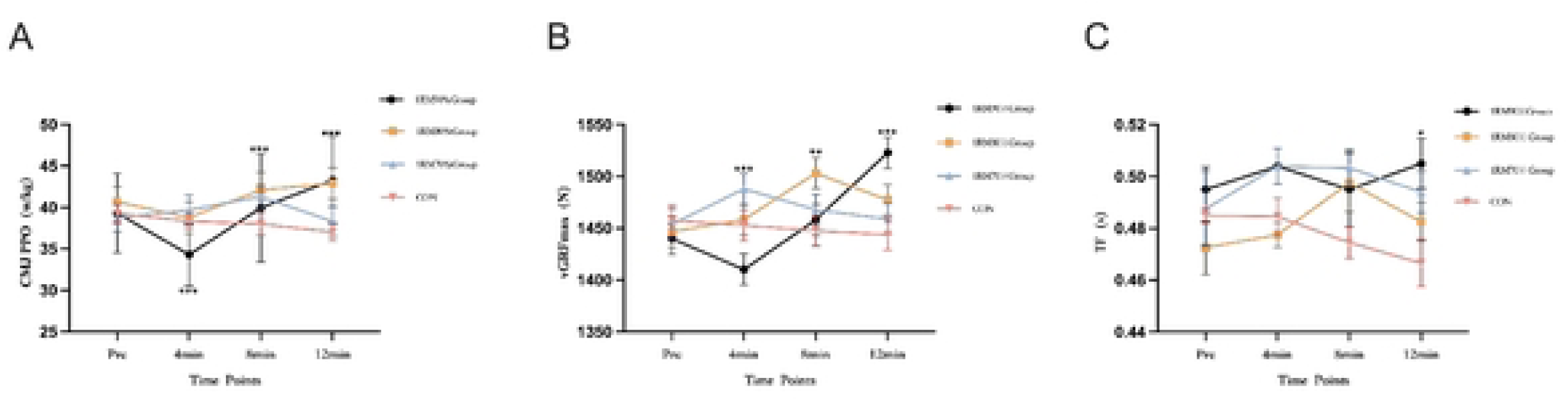
Jump kinetics during CMJ across the recovery period. A) Peak power output (PPO, W·kg⁻¹); B) Peak vertical ground reaction force (vGRF, N); C) Flight time (s). Curves show group means for the 90%1RM, 80%1RM, 70%1RM and control conditions at Baseline, 4 min, 8 min, and 12 min post-conditioning; error bars denote SD (n = 7 per group) *P<0.05,**P<0.01, *** P<0.001.

**Table 3.** Countermovement Jump (CMJ) Performance Metrics Under Different Conditioning Loads and Recovery Times (Mean ± SD)

| Metric | Group | Baseline | 4 min | 8 min | 12 min |
| --- | --- | --- | --- | --- | --- |
| PPO (W·kg) | 90%1RM | 39.33 ± 4.76 | 34.30 ± 3.76 <sup>a</sup> | 39.95 ± 6.54 | 43.30 ± 5.18 * # |
|  | 80%1RM | 40.68 ± 1.77 | 38.78 ± 1.89 # | 42.10 ± 2.15 * # | 42.90 ± 1.87 * # |
|  | 70%1RM | 38.75 ± 1.65 | 39.70 ± 1.72 # | 41.30 ± 1.31 * # | 38.30 ± 1.82 |
|  | CON | 39.25 ± 0.57 | 38.35 ± 0.48 | 38.00 ± 0.67 | 37.03 ± 0.50 |
| vGRF (N) | 90%1RM | 1439.75 ± 31.34 | 1410.00 ± 29.97 <sup>b</sup> | 1458.00 ± 59.54 | 1522.75 ± 42.03 ** |

|  |  |  |  |  | # |
| --- | --- | --- | --- | --- | --- |
|  | 80%1RM | 1446.50 ± 33.63 | 1458.25 ± 51.13 # | 1503.00 ± 40.47 ** | 1477.00 ± 37.19 |
|  |  |  |  | # |  |
|  | 70%1RM | 1453.88 ± 31.74 | 1487.75 ± 32.88 # | 1467.75 ± 35.42 | 1459.10 ± 30.15 |
|  | CON | 1457.25 ± 14.71 | 1452.75 ± 14.77 | 1447.50 ± 15.35 | 1443.75 ± 14.86 |
| FT (s) | 90%1RM | 0.49 ± 0.017 # | 0.46 ± 0.018 | 0.49 ± 0.028 | 0.50 ± 0.019 # |
|  | 80%1RM | 0.47 ± 0.002 | 0.47 ± 0.009 | 0.49 ± 0.022 # | 0.48 ± 0.015 |
|  | 70%1RM | 0.48 ± 0.028 | 0.50 ± 0.013 # | 0.50 ± 0.014 # | 0.49 ± 0.016 |
|  | CON | 0.48 ± 0.005 | 0.48 ± 0.014 | 0.47 ± 0.012 | 0.46 ± 0.018 |

**Note:** PPO = Peak Power Output; vGRF = peak vertical Ground Reaction Force; FT = Flight Time; CON = Control Group. Superscript # indicates the highest value (peak performance) within the same metric and group across time. Significance symbols indicate a significant difference compared to the CON group at the same time point: \* p < 0.05, \*\* p < 0.01. Superscript letters indicate significant differences between experimental groups at the same time point: ^a Significantly lower than the 80%1RM and 70%1RM groups; ^b Significantly lower than the 70%1RM group.

Maximal vertical ground-reaction force (vGRF). There were significant main effects of group (F(3, 24) = 7.29, p = 0.001, ηp² = 0.36) and time (F(2.25, 54.0) = 28.95, p < 0.001, ηp² = 0.65), and a significant interaction (F(6.75, 54.0) = 6.70, p < 0.001, ηp² = 0.29) (Figure 3B). At 4 min, the 70%1RM group (1487.75 ± 32.88 N) was significantly higher than the 90%1RM and control groups (p < 0.05). At 8 min, the 80%1RM group (1503.00 ± 40.47 N) showed the highest value (p < 0.01). At 12 min, the 90%1RM group (1522.75 ± 42.03 N) was highest (p < 0.01). The temporal pattern indicated that the 90%1RM group peaked at 12 min, the 80%1RM group peaked at 8 min, and control declined throughout (Table 3).

Flight time (FT). No significant main effect of group (F(3, 24) = 0.66, p = 0.593) or interaction (F(6.75, 54.0) = 2.03, p = 0.174) was observed; the main effect of time was marginal (F(2.25, 54.0) = 3.32, p = 0.065, ηp² = 0.12) (Figure 3C). Simple-effects analyses indicated significant within-group differences over time for the 90%1RM and 80%1RM groups (p = 0.036): the 90%1RM group showed a transient decrease at 4 min followed by a rebound at 12 min, whereas the 80%1RM group exhibited a slight prolongation at 8 min. Changes in the 70%1RM and control groups were not significant (Table 3).

Rectus femoris. The main effect of group was not significant (F(3, 24) = 1.22, p = 0.327, ηp² = 0.11), whereas the main effect of time was significant (F(2.25, 54.0) = 9.14, p = 0.001, ηp² = 0.43), as was the group × time interaction (F(6.75, 54.0) = 2.48, p = 0.026, ηp² = 0.38) (Figure 4A). No between-group differences were observed at baseline or 4 min (p > 0.05). At 8 min, the 90%1RM group (50.85 ± 1.50 %MVC) was significantly higher than the 80%1RM, 70%1RM, and control groups (all p < 0.05); at 12 min, the 80%1RM group (48.95 ± 6.34 %MVC) remained comparatively elevated. Simple-effects tests for the time factor indicated significant within-group changes over time in the 90%1RM and 80%1RM groups (both p < 0.05), with a marked decline in the control group (p < 0.001) (Table 4).

**Figure 4.**
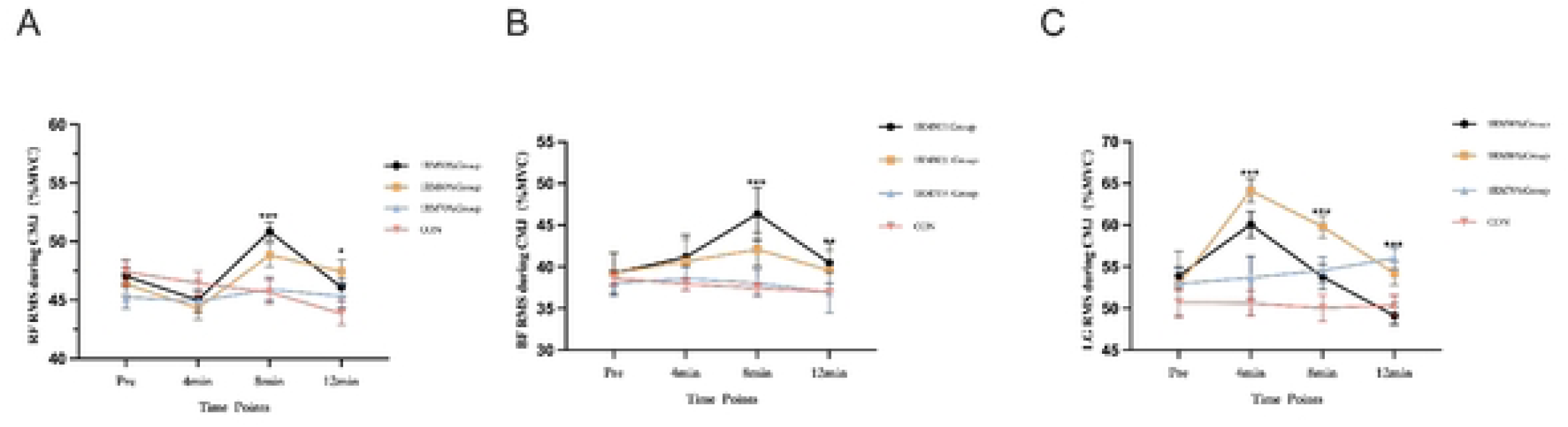
Muscle activation during CMJ.A) Rectus femoris (RF) RMS, normalized to %MVC; B) Biceps femoris (BF) RMS, %MVC; C) Lateral gastrocnemius (LG) RMS, %MVC. Traces depict the mean responses for the 90%1RM, 80%1RM, 70%1RM, and control conditions at Baseline, 4 min, 8 min, and 12 min after the conditioning squats; error bars show SD (n = 7 per group).

**Table 4.**
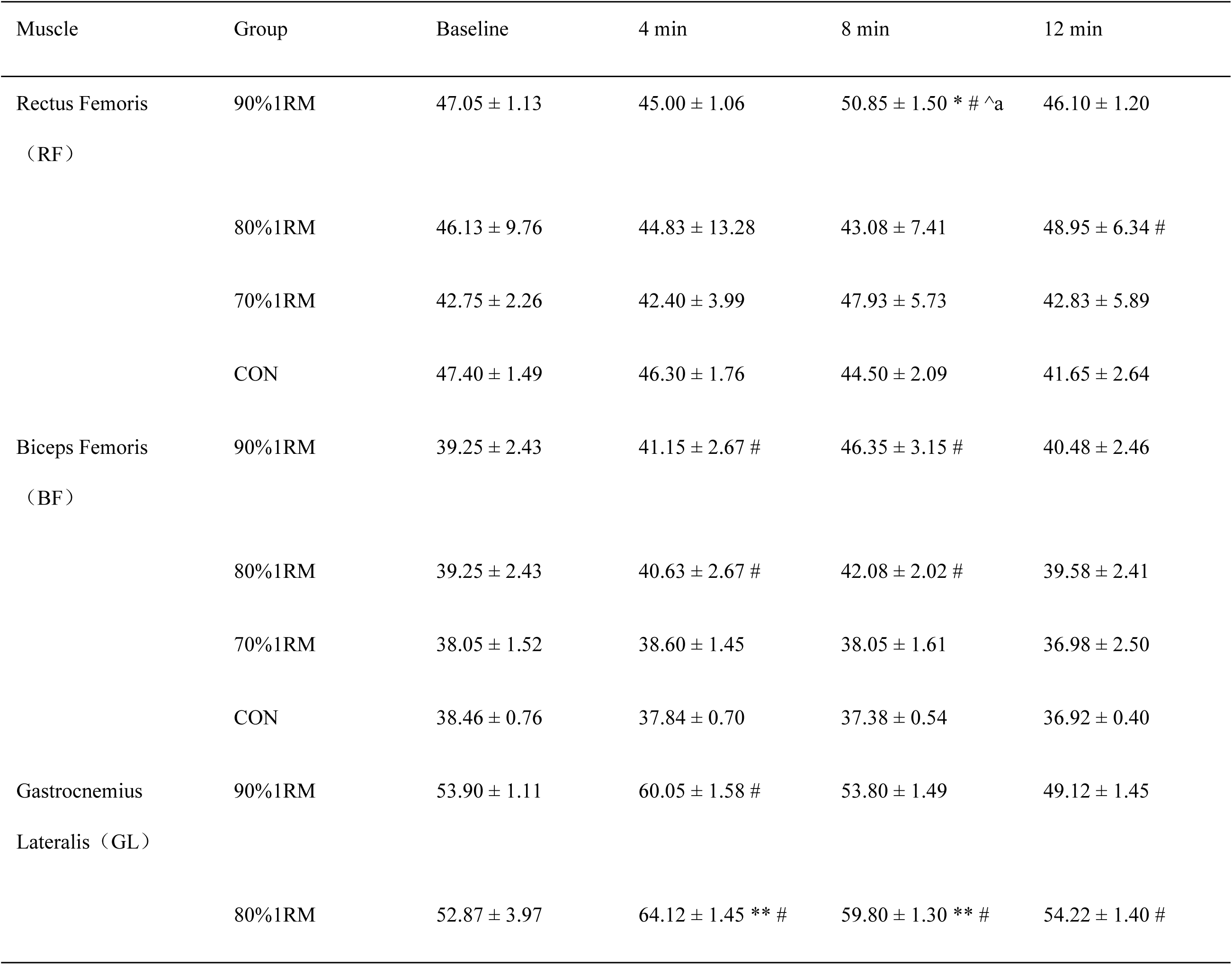
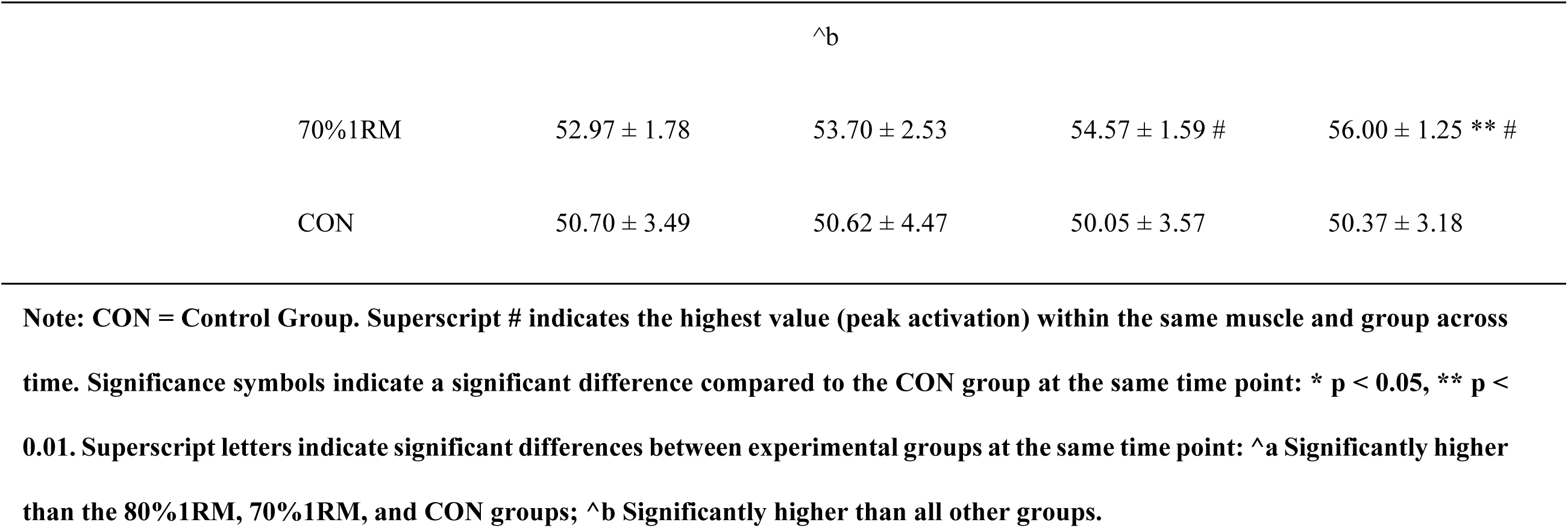
Root Mean Square (RMS) of Lower-Limb Muscles During CMJ (Mean ± SD, %MVC)

Biceps femoris. Significant main effects were found for group (F(3, 24) = 4.15, p = 0.032, ηp² = 0.15) and time (F(2.25, 54.0) = 9.14, p < 0.001, ηp² = 0.43), whereas the interaction was not significant (F(6.75, 54.0) = 1.02, p = 0.481, ηp² = 0.10) (Figure 4B). Accordingly, only main effects are reported: overall, the 90%1RM and 80%1RM groups exhibited higher biceps femoris activation than the other groups, with the control group lowest (Table 4).

Lateral gastrocnemius. The main effect of group was not significant (F(3, 24) = 0.91, p = 0.466, ηp² = 0.09), the main effect of time was significant (F(2.25, 54.0) = 37.28, p < 0.001, ηp² = 0.76), and the group × time interaction was significant (F(6.75, 54.0) = 19.36, p < 0.001, ηp² = 0.83) (Figure 4C). There were no differences at baseline (p > 0.05). At 4 min, the 80%1RM group (64.12 ± 1.45 %MVC) exceeded all other groups (p < 0.001); at 8 min, differences remained significant, with the 80%1RM group still highest (59.80 ± 1.30 %MVC, p < 0.001); at 12 min, the 70%1RM group (56.00 ± 1.25 %MVC) was highest (p < 0.01). Simple-effects tests for time showed significant within-group changes for the 90%1RM and 80%1RM groups (both p < 0.001), a gradual increase in the 70%1RM group (p = 0.044), and no significant change in control (p = 0.284) (Table 4).

## 4 Discussion

This study examined collegiate female basketball players to determine the acute effects of different back-squat loads on subsequent explosive performance and its temporal window. Overall, loads ≥80% 1RM more readily elicited post-activation performance gains. Specifically, a high load (90% 1RM) favored double-leg approach jumps at a later recovery stage (∼12 min), whereas a moderately high load (80% 1RM) produced more stable improvements at a mid-stage recovery (∼8 min). Kinetic and EMG findings corroborated each other, revealing a clear pattern in which higher loading shifted the optimal performance window further to the right in time; in contrast, performance in the control group declined naturally over time, underscoring the practical value of prescribing appropriately intense activation work in pre-competition routines.

In the double-leg approach jump (DLAJ), the 90% 1RM group exhibited the greatest improvement at the 8-min recovery time point (≈ +3.00 cm), outperforming the moderate and lower load groups overall. In the single-leg approach jump (SLAJ), the 80% 1RM group achieved the best performance at 8 min (40.50 ± 1.73 cm), significantly exceeding the control group, with minimal differences from the other loading groups. Overall, moderate-to-high squat intensities were more likely to elicit post-activation performance enhancement (PAPE), consistent with previous reviews^[30, 31]^.Canonical mechanisms include increased cross-bridge sensitivity via phosphorylation of the myosin regulatory light chain under high-intensity conditions, along with greater recruitment of high-threshold motor units (particularly type II fibers). The optimal window observed here (∼8–12 min) aligns with the commonly reported 4– 12 min range, suggesting that the time course of PAPE may be modulated by both load magnitude and individual characteristics^[32–34]^. Unlike the findings of Kaiya et al.^[35]^ in NCAA female athletes—who detected no effect when reassessing only at 5 min—our study identified significant improvements at later time points (8–12 min), likely because their sampling did not capture the highly potentiated phase that emerges after fatigue dissipates. With respect to movement specificity, we observed more stable facilitation of single-leg approach jumping at 80% 1RM. This contrasts with the report by Rogerson et al.^[36]^ in male athletes that “heavier loads yield greater effects,” implying potential sex-related differences in neuromuscular activation patterns. The overall benefit of 70% 1RM across jump modalities was limited, in line with the notion that loads below ∼75% 1RM may fail to provide sufficient mechanical tension and metabolic stress to elicit pronounced PAPE^[37]^ ^[23]^ ^[24]^。

Temporally, the 90% 1RM group peaked within 8–12 min and then declined, a trajectory consistent with the “two-factor model,” wherein performance reflects the interplay between short-term potentiation and fatigue^[38]^, The 80% 1RM group maintained a comparatively elevated level during this window, indicating a more favorable potentiation–fatigue balance. Performance in the control group declined steadily from baseline, further suggesting that, absent appropriately intense priming, neural excitability and output may naturally wane over time.

In terms of relative peak power (PPO), we observed that the 90% 1RM group peaked at 12 minutes, whereas the 80% 1RM group maintained a high and stable output across the 8–12 minute window. This finding aligns with the “force-threshold” hypothesis, which posits that the neuromuscular system must experience sufficiently intense stimulation to fully elicit the PAPE response^[39]^. Consistent with this view, Cormie et al.^[40]^ reported that lower-limb power output can be effectively enhanced when loading exceeds ∼80% 1RM. Our results closely mirror that conclusion: at 12 min the 90% 1RM group demonstrated the most robust explosive response, with both vGRF peak and PPO significantly higher than in the other loading conditions, suggesting that high-intensity loading is pivotal for eliciting comprehensive neuromuscular gains. Regarding stimulus magnitude, Villalón-Gasch et al.^[41]^ found in elite female volleyball players that heavy loading markedly improved PAPE-related performance. Similarly, Donahue et al.^[42]^ using force-platform analysis in collegiate female volleyball athletes, showed that under a high-load (NAS) protocol, 9 of 11 participants exhibited significant increases in mean propulsive force, accompanied by greater net propulsive impulse and shorter time to take-off. These findings indicate that sufficiently strong loading can produce concurrent improvements in CMJ peak power, impulse, and flight time, further underscoring the central role of robust stimuli in PAPE. Together, these results corroborate our observation that prescribing ∼80–90% 1RM with an 8–12 min recovery window can yield coordinated enhancements in PPO, vGRF, and FT.

From a recovery-time standpoint, Jensen and Ebben reported that mechanical performance gains are typically absent with recovery periods shorter than 4 minutes^[43]^, whereas McCann and Flanagan documented improvements in jump height after 4–5 minutes but did not include multidimensional kinetic indices^[44]^ 。These reports complement our use of a longer window (8–12 min), which was better suited to capturing the full expression of the PAPE effect.

By analyzing lower limb sEMG activity during the CMJ following squat loading, we found that rectus femoris activation peaked ∼8 min after the 90% 1RM condition, whereas the lateral gastrocnemius exhibited a stronger response that emerged earlier (4 min) and was sustained under the 80% 1RM condition; biceps femoris activation remained generally higher in both the 80% and 90% 1RM groups. This muscle-specific, time-dependent activation pattern aligns closely with the characteristics of PAP in female athletes described by Nuzzo^[47]^ namely, temporally distinct neural and metabolic regulation across muscles. Notably, the overall advantage observed with 80% 1RM in our study contrasts with Martinez et al.^[19]^ who reported that male athletes can achieve significant effects at ≥75% 1RM, suggesting that women’s neuromuscular systems may require a stronger external stimulus (e.g., 80% 1RM) to reach comparable motor-unit recruitment thresholds. Mechanistically, the PAPE effect induced by 80% 1RM can be viewed as the net outcome of the interplay between potentiation and fatigue^[48]^ : this intensity likely activates high-threshold motor units sufficiently while better constraining neural and metabolic fatigue, thus achieving a more favorable balance between “potentiation gain” and “fatigue cost”^[38]^ ^[30]^. This framework helps explain why rectus femoris peaked later (8 min) while the gastrocnemius responded earlier (4 min)—the optimal activation–fatigue balance point appears muscle-dependent^[38]^ ^[30]^ ^[48]^. Importantly, the performance improvements observed here were not uniformly accompanied by synchronous increases in sEMG amplitude across all muscles (e.g., biceps femoris showed an overall activation advantage rather than a pronounced time interaction), consistent with evidence that sEMG magnitude is not linearly coupled to performance; PAPE-related gains may stem more from optimized motor-unit recruitment timing than from sheer amplitude increases^[49]^. Such optimization, together with the kinetic similarity between squatting and CMJ, likely enhances the coordination of lower-limb force sequencing, ultimately translating into improvements in vertical ground-reaction force (vGRF/GRF) and rate of force development (RFD) during CMJ take-off^[50]^. Inevitably, our interpretation of sEMG signals is limited to global amplitude metrics and does not resolve activation characteristics across contraction phases, constraining insights into spatiotemporal patterns of neural drive. Future studies integrating high-resolution EMG decomposition with synchronous kinetics and muscle–tendon unit stiffness assessments may clarify how different loading schemes modulate motor-unit discharge behavior, inter-muscle coordination, and stretch–shortening cycle (SSC) efficiency, thereby refining the neuro-mechanical framework of PAPE.

## 5. Limitation

This study has several limitations: the sample size was small and drawn primarily from collegiate varsity teams, which constrains representativeness and statistical power; PAPE was assessed only at 4, 8, and 12 minutes post-intervention and did not capture earlier (≤3 min) or later (>12–30 min or even hours) response windows, potentially underestimating individuals’ optimal recovery timing and the duration of peak effects; the intervention modality was singular (back squat only), and the testing tasks focused largely on vertical jump variants, limiting generalizability across sports and movement contexts; loads and recovery intervals were not individualized based on strength level or prior responsiveness, which may have obscured true differences between stronger responders and weaker responders; recent training load, sleep– nutrition status, and menstrual cycle were not systematically controlled; and the set of monitored muscles for sEMG was limited, preventing a comprehensive depiction of lower-limb intermuscular coordination. Future research should expand and stratify the sample (including higher-performance cohorts and athletes from diverse sports), employ multiple conditioning activities (e.g., Olympic-lift derivatives, drop jumps), and implement denser and longer observation schedules; incorporate individualized prescriptions (optimizing load and recovery windows based on strength/velocity metrics); modestly increase monitoring of key lower-limb muscles and joint segments while maintaining feasibility; and record/control day-to-day factors affecting neuromuscular excitability, thereby yielding more robust, generalizable, and practice-ready evidence for PAPE in female athletes.

## 6. Conclusion

The present findings indicate that parallel back squats performed at 80–90% 1RM effectively elicit post-activation performance enhancement (PAPE). Within the 8–12 min window post-intervention, an 80% 1RM load yielded the most favorable overall benefits for single-leg jumping and multi-muscle coordinated activation, whereas a 90% 1RM load produced a more pronounced facilitation of double-leg jumping power output at the later time point (12 min). In this study, loading intensity differentially influenced neuromuscular activation and fatigue: higher loads recruited more high-threshold motor units but also imposed greater neural fatigue, thereby shifting the optimal performance window later in time. An 80% 1RM back-squat conditioning activity significantly improved basketball-specific jump performance; accordingly, integrating a moderately high load (80% 1RM) with an 8–12 min recovery period into the pre-game warm-up represents an effective strategy to enhance explosive performance in women’s basketball players.

## Ethics approval and consent to participate

The study protocol was approved by the Ethics Committee of Tianjin University of Sport (Approval No.: Tjus2025-103) and conducted in accordance with the Declaration of Helsinki. Informed consent was obtained from all participants.

## Consent for publication

Not applicable. The manuscript does not include the participant’s identification image or other personal or clinical details.

## Availability of data and materials

The data generated and analyzed during the study are provided in the main text and materials. Other related data are available from the corresponding author upon reasonable request.

## Conflict of interest

The authors declare that they have no conflict of interest.

## Funding

No external funding was received for this study.

## Authors’ contributions

SX and LYH designed the study. NSJ and ZXR performed data analysis on the collected information. SX and LYH integrated and visualized the results. SX drafted the manuscript, and LQB reviewed and revised it.

## Acknowledgements

The authors have no acknowledgments to report.

## Generative AI disclosure

This article was written by the author independently, without using any AI tools or software to generate, edit or modify the content.

## Notes

### Competing Interest Statement

The authors have declared no competing interest.

